# Female song and breeding phenology of the Nilgiri Flycatcher (*Eumyias albicaudatus*) in the Shola Sky Islands

**DOI:** 10.64898/2026.08.27.747512

**Authors:** Helly Vyas, Chiti Arvind, M Subhash, V. V. Robin

**Affiliations:** Department of Biology, Indian Institute of Science Education and Research (IISER) Tirupati, Venkatagiri Road, Tirupati, 517619, India

**Keywords:** female birdsong, BirdNET, breeding ecology, passive acoustic monitoring (PAM), sexual vocal dimorphism, Western Ghats

## Abstract

Assessing breeding phenology using singing intensity can potentially provide insights into a species’ responses to climate or habitat change. Although male passerines are known to sing elaborate songs during the breeding season, females of several tropical passerines also sing. The Nilgiri Flycatcher (*Eumyias albicaudatus*) is an endemic dimorphic songbird found in the southern Western Ghats of India. Despite its limited distribution and its ecological significance as an indicator species of cloud forests, very little is known about its vocal behaviour and breeding ecology. To examine sex-specific differences in vocalisation patterns, we analysed 294 male and 102 female focal songs from two pairs of birds. We then used data from a year-long automated recorder placed near a breeding pair to study the species’ singing phenology using the BirdNET Analyser. The two sexes broadly produced similar songs, although males’ songs included more notes at a faster pace. The sex-specific detectors failed due to this high similarity among songs of the two sexes, but we were able to automate species-level detection (F1-score = 0.795 at a confidence threshold of 0.1). We found strong seasonality in vocal activity, with peaks in singing during the breeding season, and identified a diurnal peak at dawn. More data are required to parameterise differences in annual vocal activity between the sexes. These detection patterns provide a framework for a landscape-wide examination of Nilgiri Flycatcher phenology and occurrence. This study suggests that automatic detectors can be a reliable tool for assessing singing phenology in birds and provide insights into breeding periods.

## 1. Introduction

Birds are among the most social and vocally active animals, and acoustic communication plays a crucial role in avian ecology, serving as a means for mate attraction, territorial defence, social bonding, and parent-offspring attraction (Marler, 2004; Catchpole & Slater, 2008; Podos & Webster, 2022). Bird vocalisations are correlated with behaviour and life-history events; therefore, understanding them provides insights into species’ ecology and breeding behaviour (Catchpole & Slater, 2008; Podos & Webster, 2022).

Some bird species exhibit sexually dimorphic traits, with males and females differing in plumage, body size, or ornamentation, and these differences have often been interpreted in the context of sexual selection (Price & Birch, 1996). Although bird song production has long been considered a male characteristic in songbirds (Passeriformes), studies show that female song is widespread among oscine birds and represents the ancestral state (Langmore, 1998; Odom et al., 2014). Female songs have been largely understudied due to limited documentation, a lack of sex-specific information in recordings, and misidentification of singing females as males (Webb et al., 2016; Odom & Benedict, 2018). Earlier, female songs were considered uncommon, occurring in a relatively small group of suboscines (mostly duets), in the tropical zones (Farabaugh, 1982). Recently, our knowledge of female songs has improved (Garamszegi et al., 2007; Odom & Benedict, 2018; Riebel et al., 2019), elucidating their functional role to be territorial defence (Langmore, 1998; Brunton et al., 2008; Illes, 2015), mate guarding (Rogers et al., 2007), mate attraction (Yasukawa & Searcy, 1982; Langmore et al., 1996; Morton et al., 2000), nest defence or parental care (Leonard, 2008). Studies on female bird vocalisation in Asia and South Asia have remained limited (Maller & Jones, 2001; Manshor & A., 2020; Shishkina et al., 2024). In Indian birds, studies on female songs are limited to a few species that sing regularly, such as bulbuls, mynas (Kumar, 2012), and the Oriental Magpie-Robin (*Copsychus saularis*) (Kumar & Bhatt, 2002), while others remain unexplored.

Bioacoustic technologies have emerged as powerful tools for wildlife monitoring through animals’ vocalisations (Snaddon et al., 2013; Teixeira et al., 2019). Automated recorders enable the analysis of large acoustic datasets to examine daily and annual vocal behaviour in relation to breeding phenology (Blumstein et al., 2011; Krause & Farina, 2016; Singer et al., 2025). Passive Acoustic Monitoring (PAM) is increasingly being used to monitor individual birds by their distinct vocalisations (Hutschenreiter et al., 2024; Winiarska et al., 2024; Lapp et al., 2025), and can also be used to identify sex in long-term passive acoustic monitoring of vocalisations to assess sex-specific vocal differences across a breeding cycle (Szymański et al., 2021; Appel et al., 2023; Breviglieri et al., 2026).

In this study, we examine the Nilgiri Flycatcher (*Eumyias albicaudatus*), a restricted-range bird species endemic to the Shola Sky Islands of the Western Ghats of southern India (Ali & Ripley, 1996; Somasundaram & Vijayan, 2012; Clement, 2020). The species is an insectivorous, sexually dimorphic bird, with males showing brighter blue plumage and females being duller greyish-brown (Clement, 2020). Despite being an endemic species with both sexes known to sing (Ali & Ripley, 1996), very little is known about sex-specific vocal differences in the songs of this tropical endemic oscine Passeriform, or whether such differences can be detected using algorithms on automated recordings.

In this study, we first used hand-held focal recordings to compare frequency-temporal parameters between male and female vocalisations. We then used these focal recordings to create a sex-specific classifier with BirdNET to assess sex-specific singing patterns in the Nilgiri Flycatcher across the breeding cycle. Additionally, we examined diurnal and annual changes in this species’ singing intensity over a single annual nesting cycle, using PAM and ground-truth nesting observations. We attempted to assess sex-specific differences in the singing patterns of the Nilgiri Flycatcher using an annual dataset (June 2024 - May 2025) of passive acoustic recordings from a single nesting location.

## Methods

### 2.1 Study site and sampling methods

The Nilgiri Flycatcher (hereafter NF) is endemic to the evergreen forests of the Western Ghats and is common in shola forests, which are isolated patches of jungle at high elevations (Sathiyadash et al., 2020; Sreekumar & Nameer, 2021). The breeding season for this bird is thought to span a broad window from March to June (Sathiyadash et al., 2020; Sreekumar & Nameer, 2021). This species is also known for its melodious vocalisations, which feature a complex series of 8-12 notes (Clement, 2020). We conducted our focal recording sampling in the Bombay Shola region of Kodaikanal, Tamil Nadu (longitude 77° 26’ to 77° 33’ E and latitude 10° 12’ to 10° 15’ N) between April and June 2025. Songs were sampled using a Zoom H4N handheld recorder (Zoom Corporation, 2009) with a Sennheiser ME66 shotgun microphone (Sennheiser electronic GmbH & Co. KG, 2026), with settings to record in WAV at 16-bit depth and a sampling rate of 44.1 kHz. We collected 102 songs from females (N = 2) and 294 songs from males (N = 2).

Passive Acoustic Monitoring data were collected from June 2024 to May 2025 using an autonomous acoustic recording unit, the SongMeter 4 (Wildlife Acoustics, Inc., 2024), positioned near a reused active nesting site of a banded NF pair at our field station. Nesting and fledglings have been documented in multiple years, including 2021, 2023, and 2025 (unpublished personal observations), and have been recorded as having three chicks per nesting cycle. The recorder was placed at a height of 1.5 m on a tree trunk near the nesting site of the breeding pair. The recording schedule was set to 5 minutes on and 10 minutes off (format: .wav, 16-bit depth, sampling rate: 44.1 kHz). For analysis, we filtered audio files collected between 5 am and 7 pm. During part of the same period, we conducted focal observations and recorded the banded pair’s nesting timeline, including egg laying, incubation, chick hatching, fledging, and other breeding-related behaviours (Figure 1).

**Figure 1:**
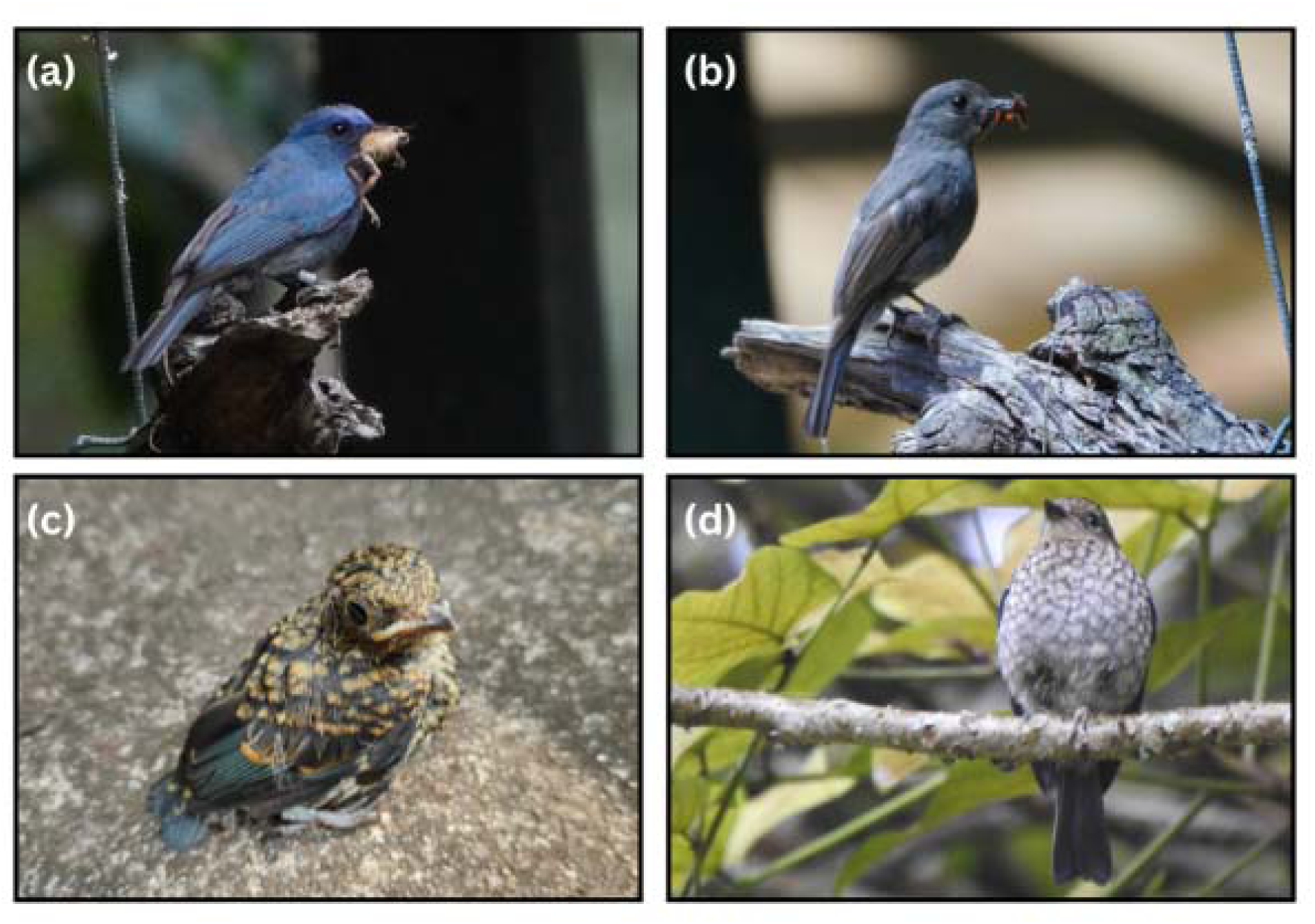
Images of a nesting a) male, b) female Nilgiri Flycatcher, both banded and with prey to feed chicks, and c) juvenile (mottled brown plumage) on the day of fledging; d) juvenile perched on a tree. (Photos a & b: Helly Vyas, c: Chiti Arvind, d: Jigyasha Khushi)

### 2.2 Focal recording analysis

Male and female songs were identified based on the singer’s plumage sexual dimorphism, as noted during recording. Only complete, high-quality recordings with a high signal-to-noise ratio were selected for analysis. All focal recordings were manually annotated at the note level in Raven Pro 1.6 (K. Lisa Yang Center for Conservation Bioacoustics, 2026), with standard settings (Hann window, a window size of 512 samples, a DFT size of 512, and 50% overlap). Each note was identified and annotated using frequency-time boxes, thus obtaining parameters of minimum frequency, maximum frequency, frequency bandwidth, mean centre frequency, peak frequency, and note duration (Figure 2). At the song level, minimum frequency, maximum frequency, frequency bandwidth, mean centre frequency, peak frequency, song duration, note count, and note pace were measured for each song. For each note, acoustic analyses were conducted at both the note and song levels. Differences between male and female songs were visualised using PCA and tested with a Wilcoxon rank-sum test, with statistical significance set at *p < 0.05*.

**Figure 2:**
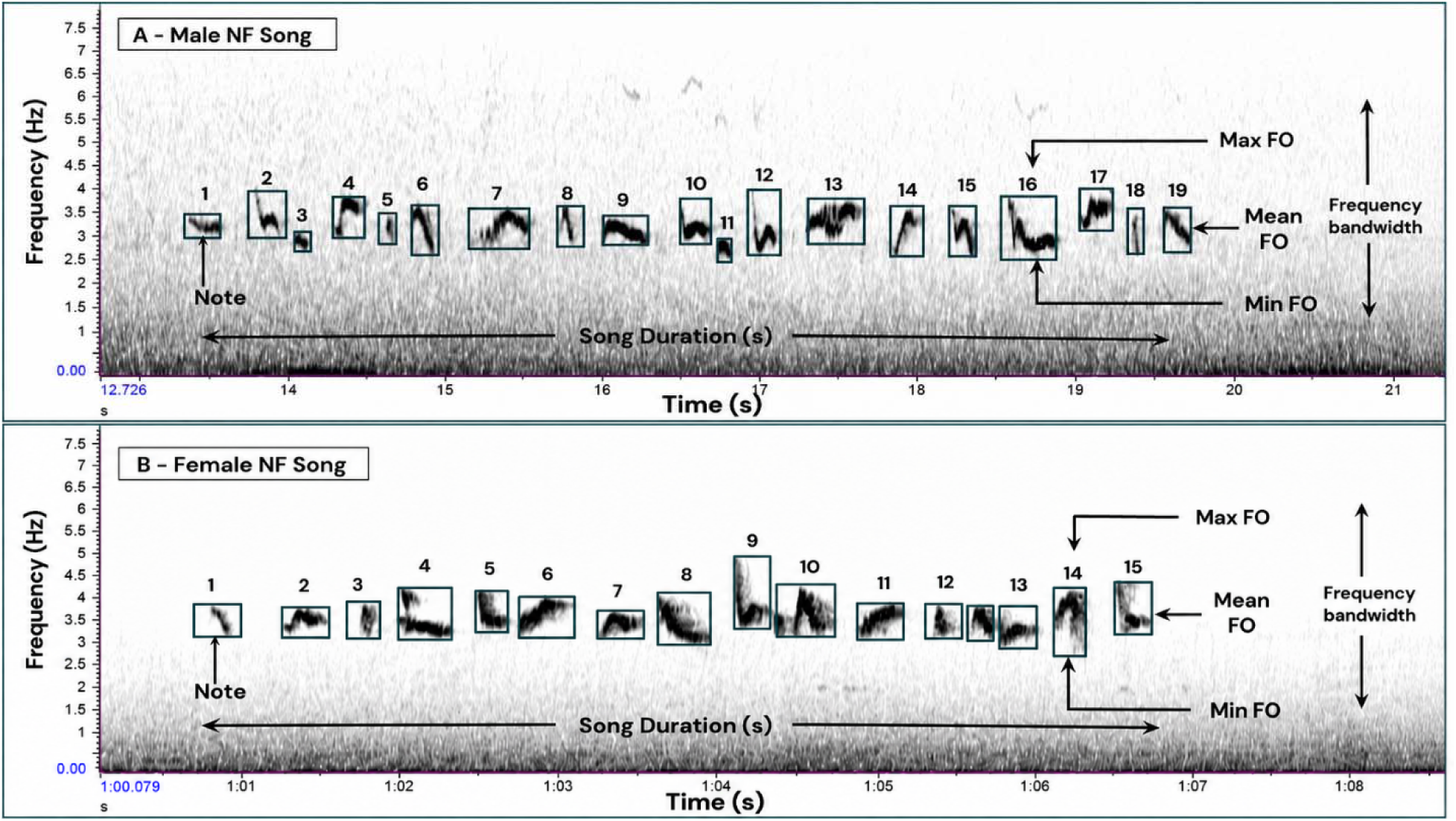
Spectrogram of a) male and b) female Nilgiri Flycatcher song with the acoustic parameters measured in the study as defined in Supplementary Table 1.

### 2.3 Passive acoustic monitoring (PAM) analysis of phenology

To investigate sex-specific songs, we developed a custom classifier to distinguish male and female NF vocalisations using BirdNET Analyser v2.4 (Kahl et al., 2021). We divided the focal data into training (80%) and validation (20%) categories. Since we did not have sex-specific truth data from the PAM itself for validation, 20% of focal segments were inserted into 5-min ARU recordings. We ensured that recordings used for training were not included in the validation set. For training, we had three target classes: male and female voices, and environmental sounds (e.g., rain, insects, wind, and other background noises) as a noise class. We validated the performance of the sex-specific classifier by manually validating 1000 segments with confidence scores ranging from 0.1 to 1.0. We assessed the confidence threshold by fitting regression curves to male and female vocalisations. Performance was assessed for the male and female classes using precision, recall, and F1-score, with overall classification accuracy evaluated across confidence thresholds (0.1 to 1.0).

As the performance of the sex specific classifier was not robust (discussed below), we decided to merge the sexes for detection throughout the year. This strategy would not yield sex-specific patterns, but it would still provide species-level patterns. We used a custom species list that included only the NF in the BirdNET Analyser to automatically detect the targeted species’ vocalisation. BirdNET was run with the default parameter settings (confidence score threshold = 0.1, sensitivity = 1, overlap = 0, audio speed modification = 1, minimum bandpass frequency = 2000 Hz, maximum bandpass frequency = 4500 Hz, and generation of 6-s segments for species detection and it attributes a confidence score to each detection, ranging from 0.1 to 1. We validated 1200 segments (segment length = 6 sec; confidence score > 0.1) from the winter and summer months to generate the logistic regression curve for the default BirdNET classifier.

Based on previous breeding records (Ali & Ripley, 1996; Somasundaram & Vijayan, 2012), we categorised the months into breeding (March to June) and non-breeding (July to February). To evaluate the performance of the default BirdNET classifier using precision, recall, and F1 score (Knight et al., 2017; Fairbairn et al., 2025), we selected validation days from the breeding (March-May 2025, 935 minutes) and non-breeding (October-December 2024, 350 minutes) periods. Default classifier performance was evaluated across confidence thresholds of 0.1 to 1.0 for both the breeding and non-breeding periods.

### 2.4 Diel and annual vocal activity analysis

Using the BirdNET classifier’s output, we analysed daily acoustic activity of NF across the year. We tested the temporal distribution of annual and diel vocal activity using circular statistics (circular package; Agostinelli) (Agostinelli & Lund, 2025). Mean direction (μ), mean vector length (r), and circular standard deviation (SD) were generated to describe the timing and concentration of diel and annual vocal activity. Rayleigh’s test was used to test whether vocal activity was uniformly distributed across time.

## Results

### 3.1 Sex-specific vocal differences in acoustic parameters

During the breeding season (May-June 2025), we recorded female (n=2 individuals; n=102 songs) and male (n=2 individuals; n=294 songs) NF and extracted multiple acoustic parameters for analysis. Significant differences were observed between male and female songs and notes for most acoustic parameters (Figure 3, Supplementary Table 2, and Supplementary Figure 1).

**Figure 3:**
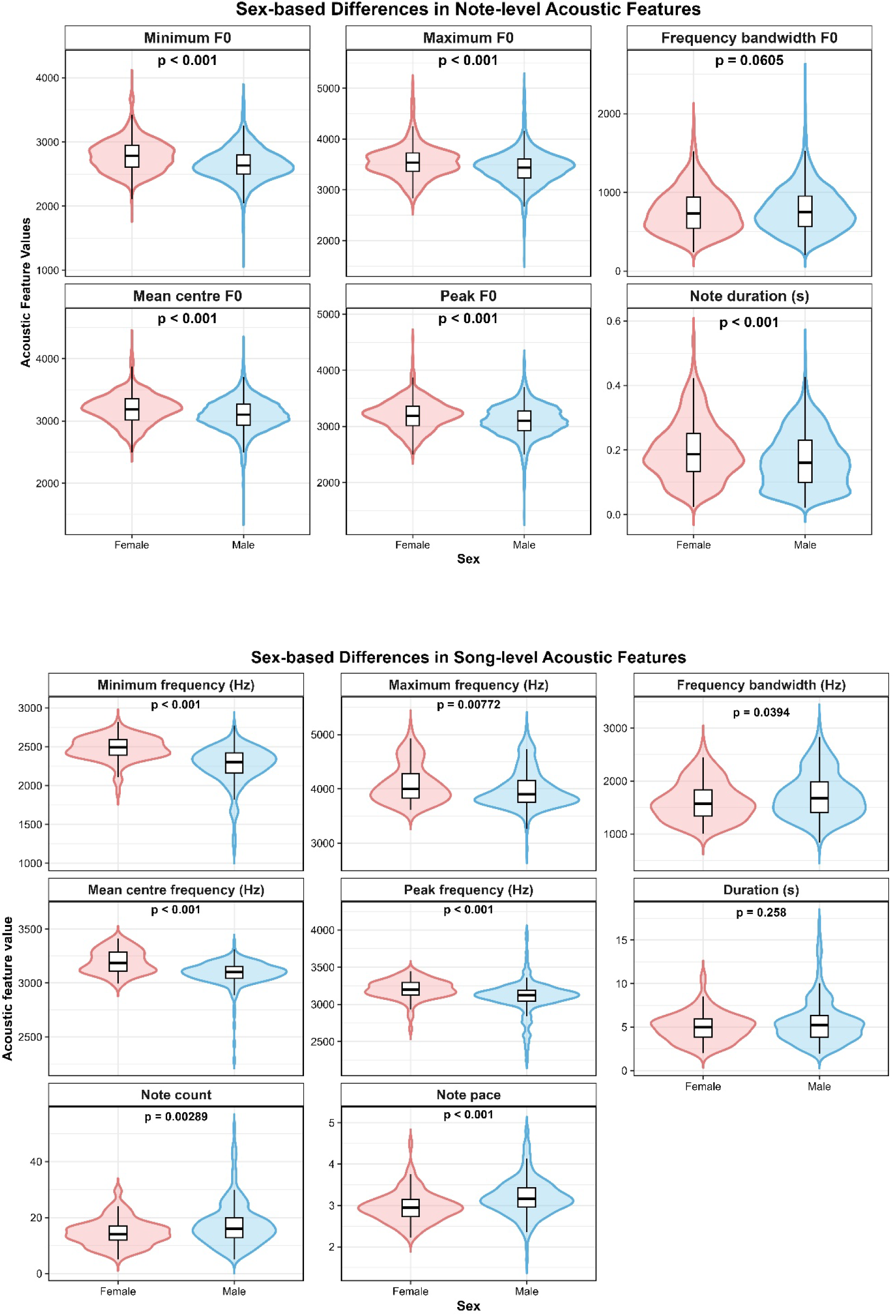
Comparison of note-level (above) and song-level (below) acoustic parameters between male and female Nilgiri Flycatcher (Eumyias albicaudatus) using violin plots. At the note level, we observed significant differences in the frequencies of minimum, maximum, mean centre, peak, and note duration, while frequency bandwidth did not differ significantly between sexes. At the song level, females had higher minimum, maximum, mean centre, and peak frequencies than males, while males had greater frequency bandwidth, note count, and note pace. Song duration did not differ significantly between sexes.

At the note level, we find differences in the frequency-level parameters between males and females (minimum frequency, maximum frequency, mean centre frequency, and peak frequency; all p < 0.001, Supplementary Table 2). Female notes were also significantly longer in duration than male notes (p < 0.001). Song duration did not differ significantly between the sexes (Male = 5.54 s ± 0.17, Female = 5 s ± 0.17, *p = 0.258*) but males had a significantly larger number of notes per song (Male=17.52 ± 0.51, Female=14.69 ± 0.50, *p = 0.0028*) and subsequently greater note pace (Male = 3.20 ± 0.03, Female = 2.97 ± 0.04, p = 7.592 × 10 ). Females produced songs with higher frequency characteristics, including higher minimum frequency (Female = 2479.72 ± 15.41 Hz, Male = 2261.60, *p* < 0.001), maximum frequency (Female = 4102.48 ± 36.09 Hz, Male = 3995.36, *p* = 0.0077), mean centre frequency (Female = 3197.70 ± 10.99 Hz Male = 3091.06 ± 7.56 Hz, *p* < 0.001), and peak frequency (Female = 3204.99 ± 13.21 Hz, Male = 3099.02 ± 14.28 Hz, *p* < 0.001) than males. Males also showed a significantly greater frequency bandwidth than females (Male = 1733.76 Hz ± 28.26, Female = 1622.76 Hz ± 37.27, *p = 0.039*) (Supplementary Table 2).

### 3.2 Evaluation of the sex-specific classifier and BirdNET species-level classifier

Evaluating the sex-specific custom classifier using regression curves (Supplementary Figure 7), we found that performance was poor for both males (precision = 0.52, recall = 0.50, and F1-score = 0.05) and females (precision = 0.51, recall = 0.50, and F1-score = 0.03), with an overall accuracy of 0.04. Low accuracy might be due to limited training data for both sexes, which may be insufficient to differentiate between them and may explain the classifier’s poor performance.

On the other hand, the species-level classifier (BirdNET default) performed well, and the logistic curves indicated a positive correlation between BirdNET confidence scores and true detection (Supplementary Figure 5). We chose a threshold of 0.1 for further analyses to maximise detections and include as many true positives as possible while maintaining high precision. At the 0.1 confidence threshold, performance metrics were the highest (precision = 0.84, recall = 0.77, and F1-score = 0.79; Supplementary Figure 6)

### 3.3 Diurnal pattern and annual vocal activity patterns using BirdNET detections

Using the annual passive acoustic dataset from June 2024 to May 2025, we compared diurnal vocal activity between the breeding (March-June) and non-breeding (July-Feb) periods. We found that both periods showed distinct diurnal variation (Figure 4). During both breeding and non-breeding seasons, diurnal vocal activity is highest in the mornings, peaking shortly after sunrise (06:00 to 09:00 AM), with significantly higher activity in the breeding season (237 ± 127 mean diurnal detections/day) than in the non-breeding season (142 ± 59.4 mean daily detections/day, W = 6121.5, p < 0.001). Differences in hourly vocal activity between breeding and non-breeding periods were especially pronounced at 1100hrs, with vocal activity four times higher (average detections/hour: breeding season 11.56 ± 1.62; non-breeding 2.50 ± 0.41). At 0800hrs, vocal activity was similar throughout the year (breeding 22.61 ± 2.23 vs. non-breeding 20.83 ± 2.67 avg detections/hour). Vocal activity at 0600 hrs is the highest year-round, with twice as much activity in the breeding period (31.34 ± 3.10 avg detections/hour) as compared to the non-breeding period (11.62 ± 1.24 avg detections/hour) (Figure 4). Circular statistics further showed that vocal activity was not evenly distributed throughout the day but was higher around particular times (mean direction = 0.812, *r* = 0.202, circular SD = 1.788, Rayleigh test = 0.202, *p* < 0.001; Supplementary Table 3).

**Figure 4:**
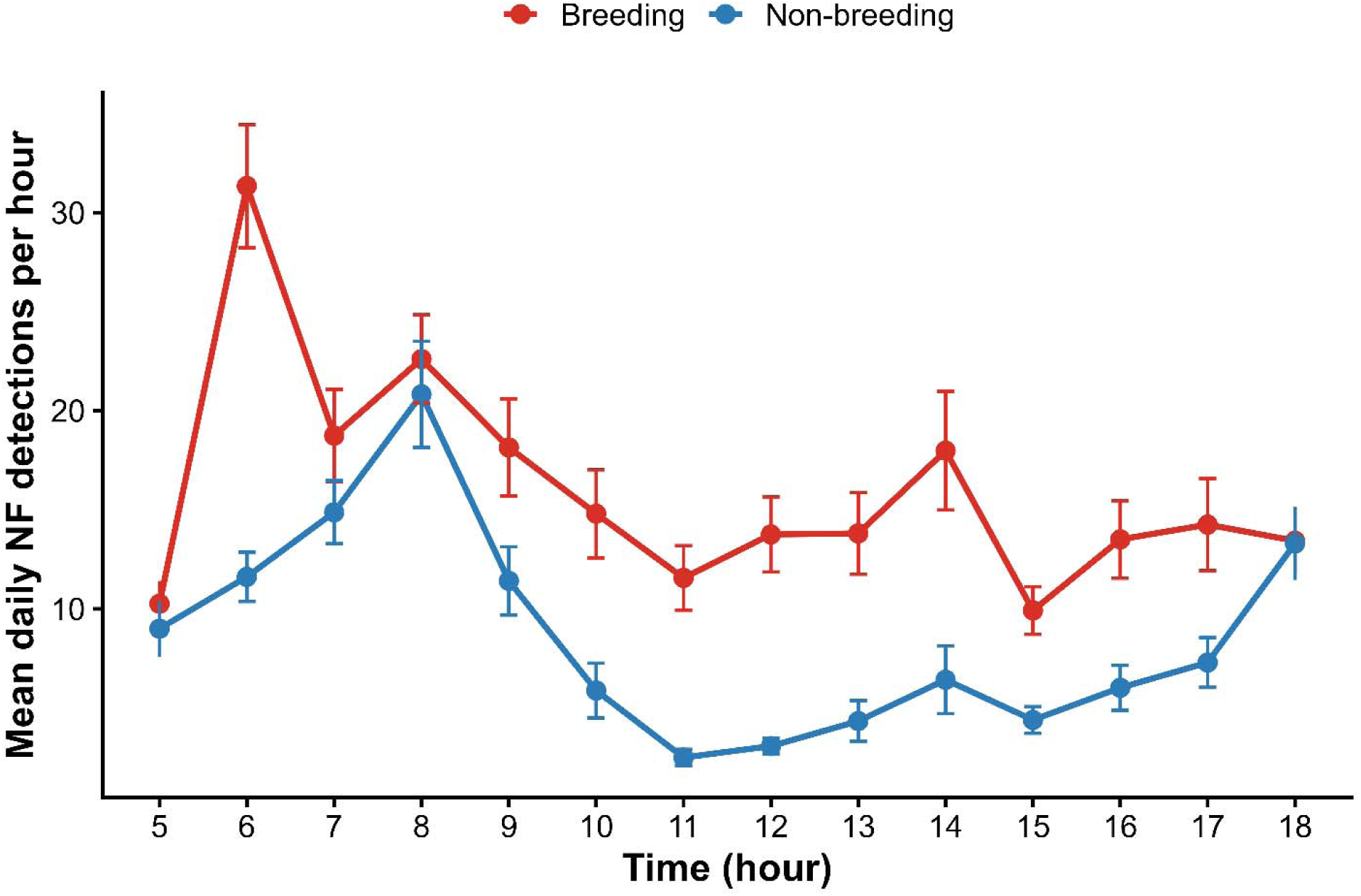
Average hourly (24hr format) BirdNET detections of the Nilgiri Flycatcher during the breeding (March-June) and non-breeding (July-Feb) periods. Values represent the mean number of BirdNET detections per hour across recording days. Error bars represent the standard error of the mean (SE).

For the annual dataset (June 2024-May 2025), we find a clear seasonal variation in the NF’s vocal activity (Figure 5), with peak mean daily detections in March (248 ± 137 detections per day; Supplementary Figure 8). This coincided with our observations of the first nesting event with egg-laying (8th March) and hatching (26 March). A second nesting attempt was recorded at the end of May, when two eggs were found in the same nest, indicating that a pair can have more than one breeding attempt in a season. Circular statistics further showed that annual vocal activity was significantly seasonal (r = 0.590, p < 0.001; Supplementary Table 3), indicating that vocal activity was greater, particularly during the breeding period (Feb to May), rather than uniformly distributed across months (see Supplementary Figure 8).

**Figure 5:**
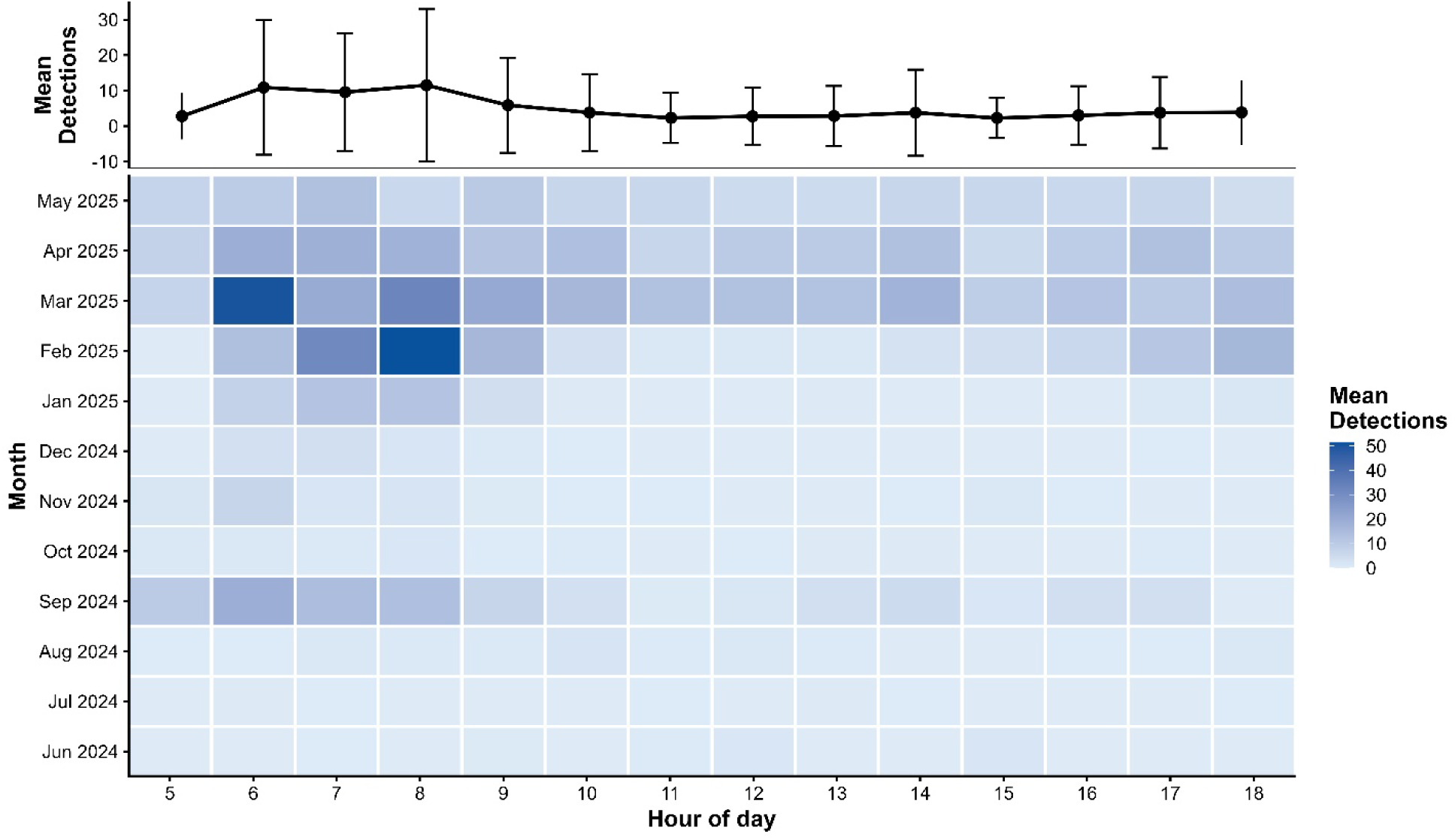
The heatmap shows the mean number of Nilgiri Flycatcher detections per day for each month and hour. The line plot above shows mean detections across hours over the annual recording period for the passive acoustic dataset (June 2024 to May 2025). The study period included both the breeding (March-June) and non-breeding (July-Feb) periods. Colour intensity represents the mean number of BirdNET detections recorded during each hour from 05:00 to 18:00 h across different months.

## 4. Discussion

This study extends our knowledge of female vocalisations in oscine Passeriformes; we find significant differences between male and female songs of the NF and subsequently use PAM to monitor breeding activity.

In several tropical species, we find sex-specific differences in vocalisations that serve different functional roles (Mennill & Vehrencamp, 2005; Odom et al., 2021; Liu et al., 2025). Adding to the growing body of tropical female bird songs, we characterise differences and similarities between male and female songs of the NF, endemic to the WG biodiversity hotspot in India. Although breeding-period songs are not aurally discernible between males and females, we do find spectral differences, with males’ songs being at a lower frequency than female songs. Similar sex-specific differences in song frequency have been reported in some species, which might be related to social functions such as territorial defence, mate guarding, or sex-specific response (Mennill & Vehrencamp, 2005; Odom et al., 2016; Fishbein et al., 2018). In the Nilgiri flycatcher, notably, the songs of both sexes are comparable in length, but males sing more notes, resulting in a higher note rate than females. Note-pace differences between sexes (Mennill & Vehrencamp, 2005; Odom et al., 2021) may be associated with breeding synchrony, sex-specific vocalisations, functional roles such as territorial defence, or reproductive success (Cain et al., 2015; Brunton et al., 2016; Odom et al., 2021). An increase in females’ overall song rate is associated with reproductive success (Cain et al., 2015; Brunton et al., 2016). In a similar vein, we did notice females singing more during the feeding/provisioning phase than males, although this remains to be quantified (personal observation). Very little is known about the function of female songs, but recent studies suggest that female songs during the breeding season also have a functional role in tutoring the young ones (Riebel, 2016; Evans & Kleindorfer, 2016). Stereotypically, male birds learn from male tutors (Böhner, 1983; Hausberger et al., 1995; Wheelwright et al., 2008; Mooney, 2022). As previous studies show that both sexes sing, but song learning is sex-specific in species (Wickler & Seibt, 1988; Jordan Price, 1998; George & Cousillas, 2013), which can often be same sex tutors to same sex nestlings and, in some species, both sexes contribute equally to being tutors (Geberzahn & Gahr, 2013; Evans & Kleindorfer, 2016).

Using annual automated recording data, we find strong seasonal patterns in NF vocalisations. Singing intensity increases as the breeding period approaches (Best, 1981; Chen et al., 2015; Yuzyk & Yuzyk, 2025), indicating that increased singing serves functional roles in mate attraction, parental care, and pair-bond strengthening during the breeding season (Nowicki & Searcy, 2005; Cain & Langmore, 2015). Daily patterns indicated more songs in the morning, irrespective of breeding, coinciding with the dawn chorus reported in tropical birds under light conditions at dawn, suggesting a correlation between the timing of dawn song and minor changes in photoperiod (Berg et al., 2006; Quispe et al., 2017). However, we found that during the breeding period, birds are more vocally active overall in the afternoon than in the non-breeding period; for example, at 11:00, birds vocalised approximately 4 times as much as in the non-breeding period. Vocal activity of NF remains higher at 08:00 during both breeding and non-breeding periods. Similar variation in diurnal and annual singing patterns has been observed in other species (Moran et al., 2019; Szymański et al., 2021), suggesting a potential marker period for targeting PAM recordings and identifying breeding instances by time of day. Song rate later in the morning – around 11 am, could be used as a sign of breeding, while earlier in the morning (6 am to 8 am) could be used to detect presence. This can be useful for future survey design of species in protected areas through PAM (Clapp et al., 2026). Male and female vocal activity may also vary over temporal scales, with sex-specific daily and annual vocal patterns (Patchett et al., 2021; Szymański et al., 2021). Although our custom sex-specific classifier was unsuccessful, perhaps due to the small differences between male and female songs and limited sample sizes, we emphasise the need for better sex-specific training classes to leverage PAM datasets and tease apart the roles of male and female songs.

Overall, through focal recordings, we find vocal sexual dimorphism in the Nilgiri Flycatcher, a visually dimorphic endemic tropical bird. This species also displays strong vocal seasonality, peaking during the nesting period, corroborated by visual observations. We suggest that Passive Acoustic Monitoring datasets, paired with analytical ML tools such as the BirdNET Analyzer, are useful for long-term species monitoring in protected areas.

## 6. CRediT authorship contribution statement

HV - data curation, analysis, investigation, methodology, visualisation, writing (original draft and review); CA - conceptualisation, data curation, funding acquisition, investigation, methodology, project administration, writing (original draft and review); MS - investigation; VVR - conceptualisation, funding acquisition, resources, supervision, writing (review).

## 8. Declaration of Artificial Intelligence usage

Grammarly was used to check grammar, spelling, and language. Claude was used for literature search assistance, and ChatGPT was used to assist with R code, including debugging and troubleshooting errors. All content has been reviewed and revised by the authors. The authors take full responsibility for the accuracy and originality of the work.

## 9. Acknowledgements

We thank the Tamil Nadu Forest Department for granting the necessary permits (Proc. No.: WL5(A)/38623/2020; Permission No.: 46/2023) to conduct this study. We are grateful to the field team for their help with observational and acoustic data collection. We thank Viral Joshi, Kunal Jagetiya, and Aditya Panigrahy for their guidance with the BirdNET analyses. We thank Viral Joshi, Vinay KL, and members of the Bird Ecology Lab at the Indian Institute of Science Education and Research, Tirupati, for feedback on the manuscript. We also thank the Kodaikanal International School for providing accommodation during the fieldwork.

## 10. Data availability

All code and data will be available on request.

