## Supplementary Material for "Female song and breeding phenology of the Nilgiri Flycatcher (*Eumyias albicaudatus*) in the Shola Sky Islands"

**Supplementary results:**

**Supplementary Table 1. List of measured call features.**

| **Acoustic measures** | **Definition** |
| --- | --- |
| Minimum F0 | Minimum frequency of the fundamental frequency (F0). |
| Maximum F0 | Maximum frequency of the fundamental frequency (F0). |
| Delta F0 | Difference between the maximum and minimum fundamental frequency (Maximum F0 - Minimum F0). |
| Mean center F0 | Mean value of the fundamental frequency across the vocalisation. |
| Peak F0 | Fundamental frequency at which the maximum energy or amplitude occurs. |
| Duration (S) | Total length of the vocalisation, measured from the beginning to the end of the song. |
| Note count | Total number of individual notes or syllables within a vocalisation. |
| Note pace | The rate at which notes are produced, calculated as the number of notes per total duration of vocalisation. |

**Supplementary Table 2:  Comparisons of acoustic parameters of songs and notes between male and female Nilgiri Flycatcher (*Eumyias albicaudatus*)**

| **Acoustic parameter** | **Males** | **Females** | **Test statistic (W)** | **p value** |
| --- | --- | --- | --- | --- |
| **Song-level parameters** | | | | |
| **Minimum frequency** | 2261.6 ± 16.16 | 2479.72 ± 15.41 | 18160.5 | <0.001 |
| **Maximum frequency** | 3995.36 ± 23.28 | 4102.48 ± 36.09 | 13478.0 | 0.0077 |
| **Frequency bandwidth** | 1733.76 ± 28.26 | 1622.76 ± 37.27 | 9735.5 | 0.0393 |
| **Mean centre frequency** | 3091.06 ± 7.56 | 3197.70 ± 10.99 | 16990.0 | <0.001 |
| **Peak frequency** | 3099.02 ± 14.28 | 3204.99 ± 13.21 | 16025.0 | <0.001 |
| **Duration (s)** | 5.54 ± 0.17 | 5.00 ± 0.17 | 10472.0 | 0.2581 |
| **Note count** | 17.52 ± 0.51 | 14.69 ± 0.50 | 9011.5 | p = 0.0029 |
| **Note pace** | 3.20 ± 0.03 | 2.97 ± 0.04 | 7451.0 | <0.001 |
| **Note-level parameters** | | | | |
| **Minimum F0** | 2652.80 ± 4.07 | 2786.84 ± 6.58 | 3,935,915 | <0.001 |
| **Maximum F0** | 3431.81 ± 5.03 | 3549.38 ± 8.28 | 3,682,647 | <0.001 |
| **Frequency bandwidth F0** | 779.01 ± 4.50 | 762.54 ± 7.35 | 2,938,418 | 0.0605 |
| **Mean centre F0** | 3093.89 ± 3.99 | 3196.27 ± 6.21 | 3,752,698 | <0.001 |
| **Peak F0** | 3106.67 ± 4.24 | 3208.33 ± 6.70 | 3,683,739 | <0.001 |
| **Note duration (s)** | 0.17 ± 0.00 | 0.20 ± 0.00 | 3,585,600 | <0.001 |

**Supplementary Table 3:** Circular Statistics for annual and diel variation of Nilgiri Flycatcher vocal activity.

| **Analysis** | **Mean direction** | **Mean vector length (r)** | **Circular SD** | **Rayleigh test statistic** | **P-value** |
| --- | --- | --- | --- | --- | --- |
| **Annual vocal activity** | 5.026 | 0.590 | 1.027 | 0.590 | < 0.001 |
| **Diurnal vocal activity** | 0.952 | 0.342 | 1.464 | 0.342 | < 0.001 |

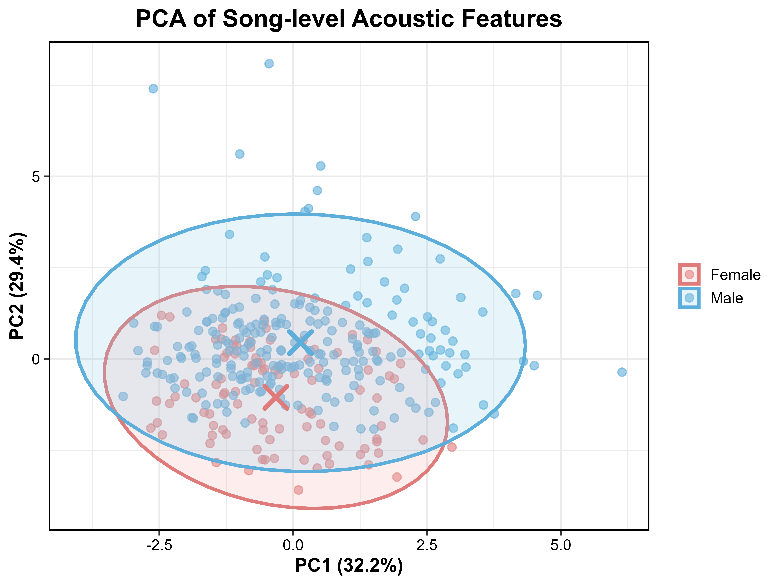

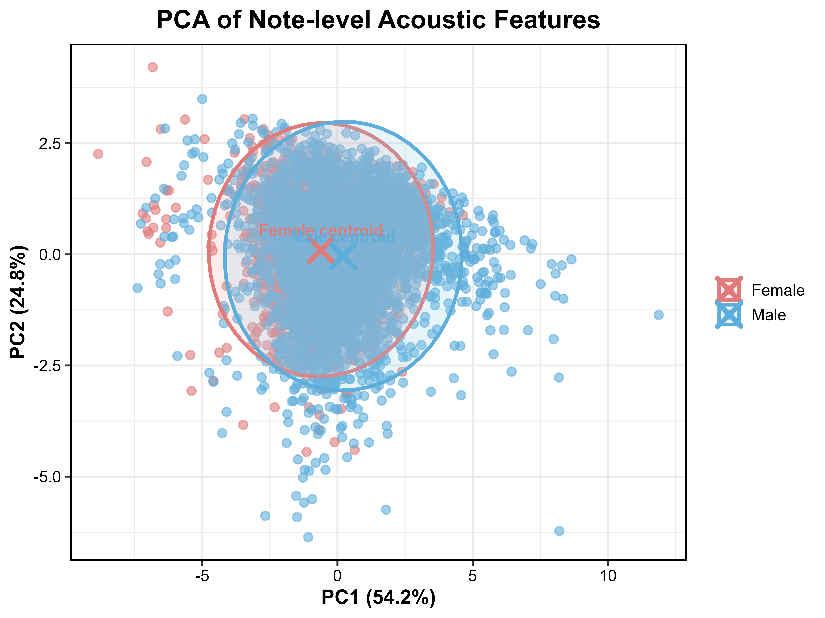

***Supplementary Figure 1: A:*** *PCA plot indicates no distinct separation between male and female vocalisations at the note level, but some differences at the song level.* ***B:*** *Each point represents an individual note in Supplementary Figure 1 (a) and a song in Figure 1 (b), while shaded ellipses indicate 95% confidence intervals for each sex. PC1 is mainly associated with frequency bandwidth, note count, and song duration, while PC2 is primarily associated with mean centre frequency, lowest frequency, peak frequency, and note pace (Supplementary Figure 2).*

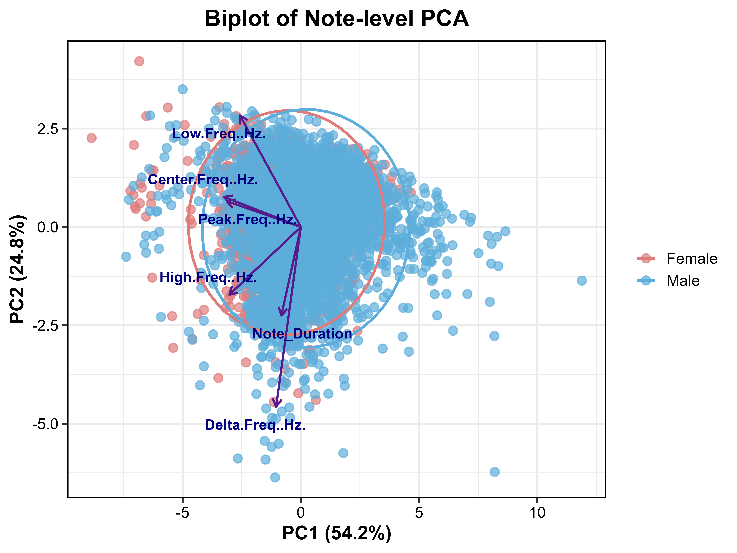

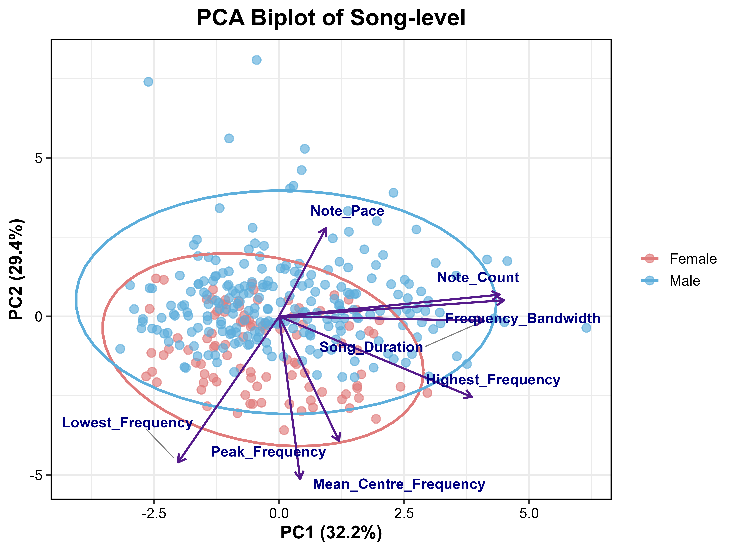

***Supplementary Figure 2:*** *Principal Component Analysis (PCA) biplots of (a) note-level and (b) song-level acoustic features of male and female Nilgiri Flycatchers (Eumyias albicaudatus).*

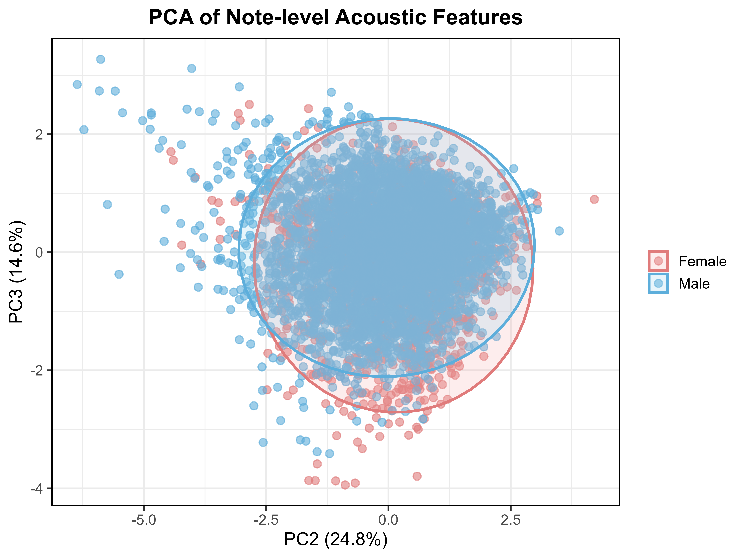

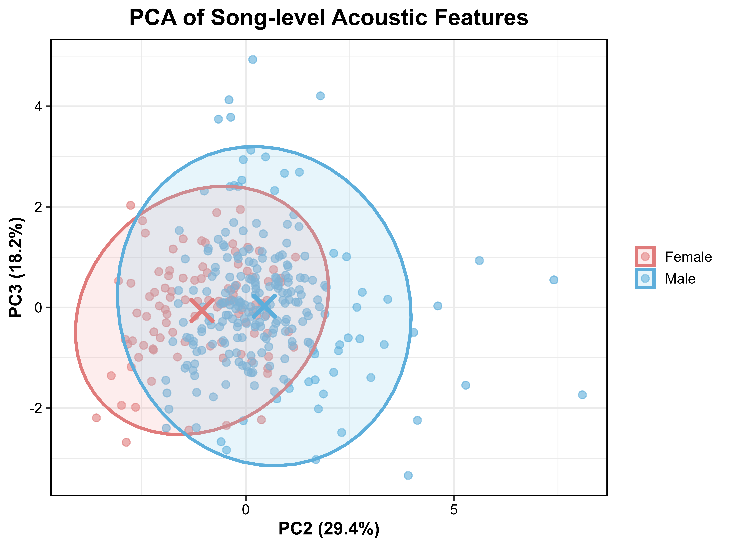

***Supplementary Figure 3:*** *PCA score plots of acoustic features in male and female Nilgiri Flycatchers (Eumyias albicaudatus): (a) note-level acoustic features (PC2 vs. PC3); and (b) song-level acoustic features (PC2 vs. PC3).*

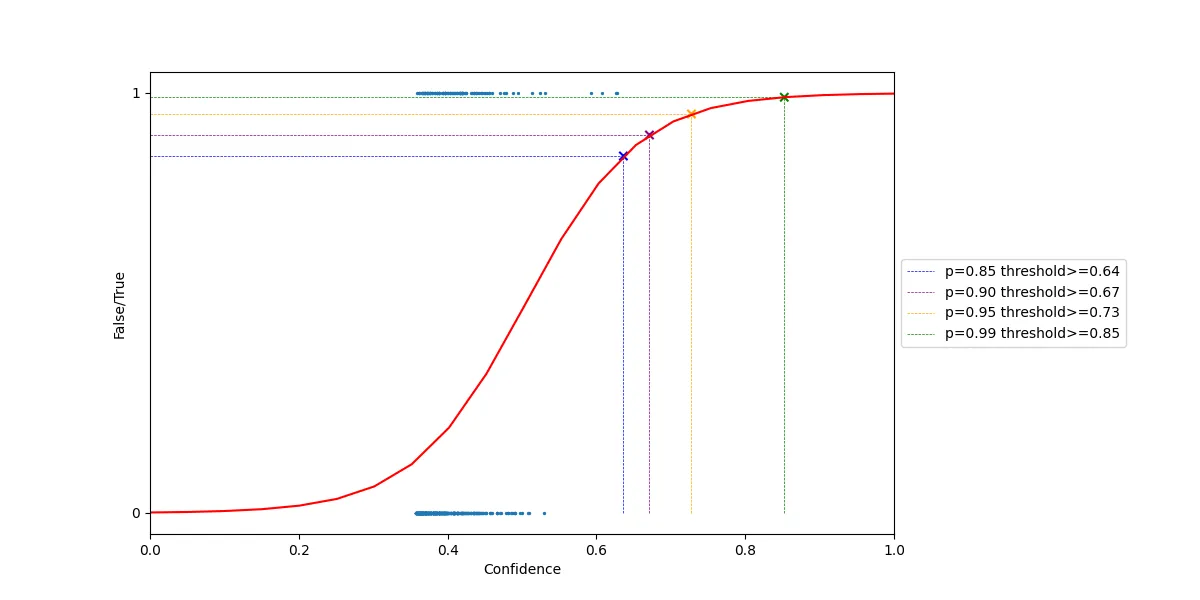

***Supplementary Figure 4 (a):*** *Regression curve for the female custom classifier*

*Figure summary - The figure shows the regression curve for the female customer classifier. The red curve represents the relationship between classifier confidence (x-axis) and the false/true detections (y-axis). The classifier's performance increases with confidence, approaching 1 at higher confidence values.*

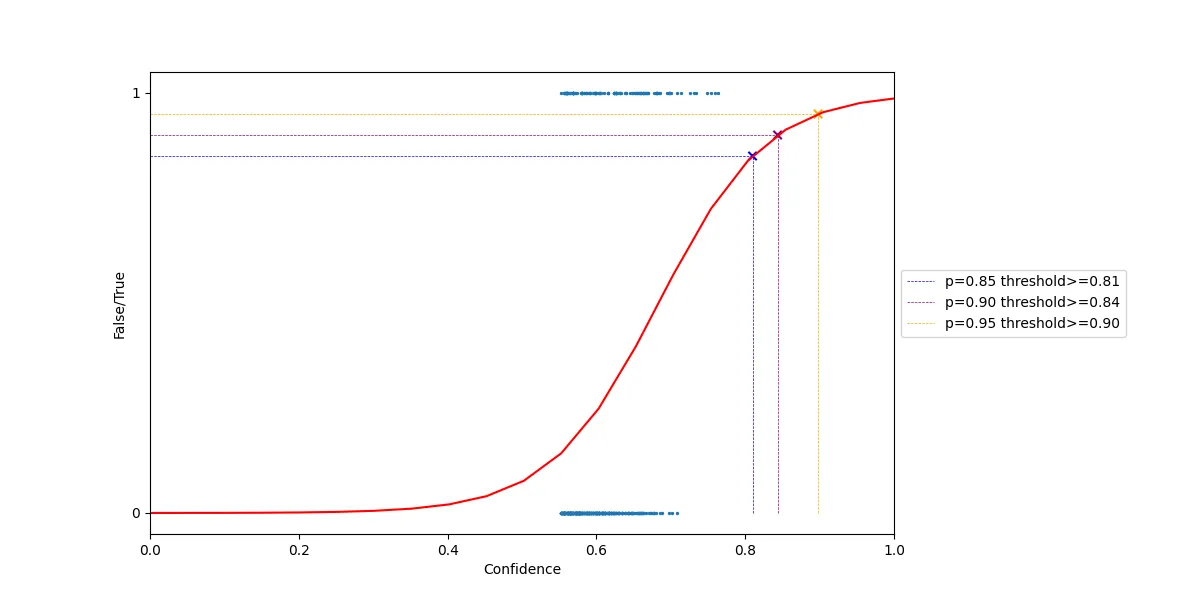

***Supplementary Figure 4 (b):*** *Regression curve for the male custom classifier*

Figure summary - The figure shows the regression curve for the male custom classifier. The red curve shows the relationship between classifier confidence (x-axis) and the False/True detections (y-axis). The curve increases between 0.6 and 0.9 confidence, showing improved classification performance at higher confidence values.

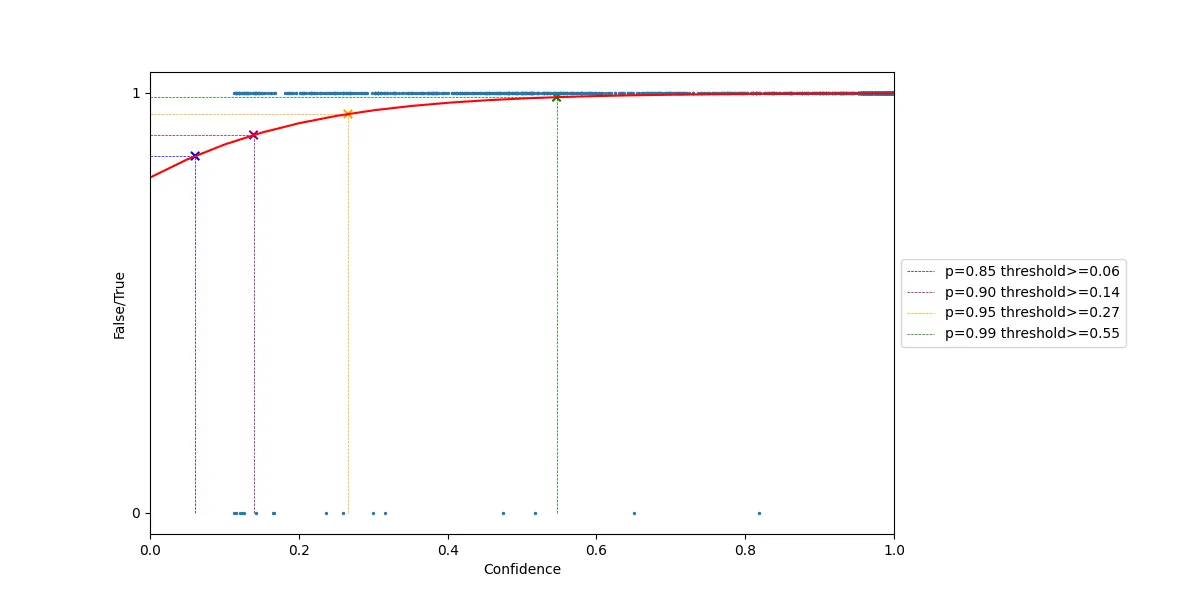
***Supplementary Figure 5:*** *Logistic regression curve showing the relationship between BirdNET confidence scores and the probability of a true detection for Nilgiri Flycatcher (Eumyias albicaudatus).*

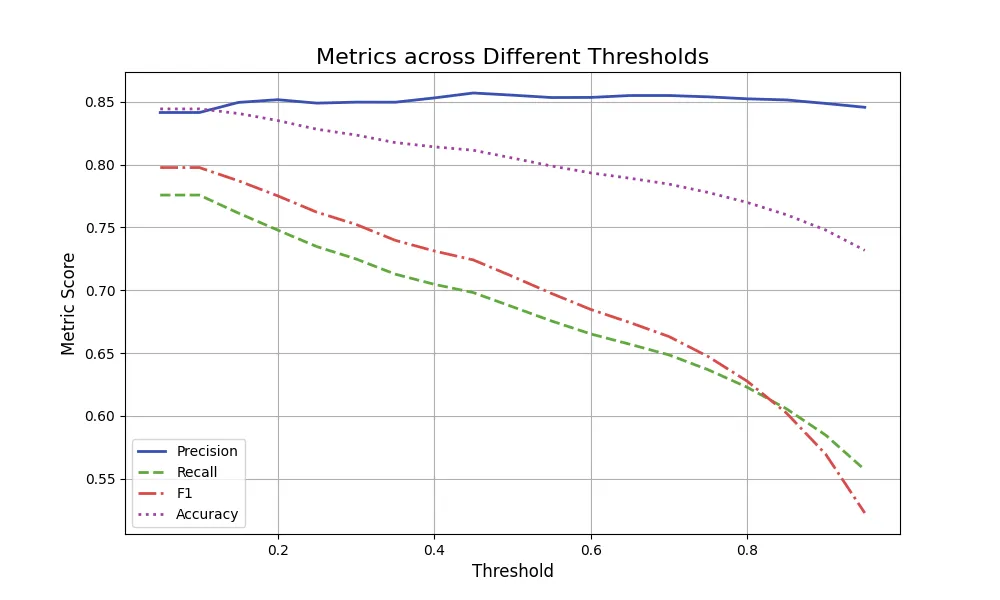

***Supplementary Figure 6:*** *Performance of the default BirdNET classifier across different confidence thresholds for detecting Nilgiri Flycatcher vocalisations during the breeding and non-breeding periods* (6 months)*. Precision, recall, F1-score, and accuracy are shown as a function of the confidence threshold.*

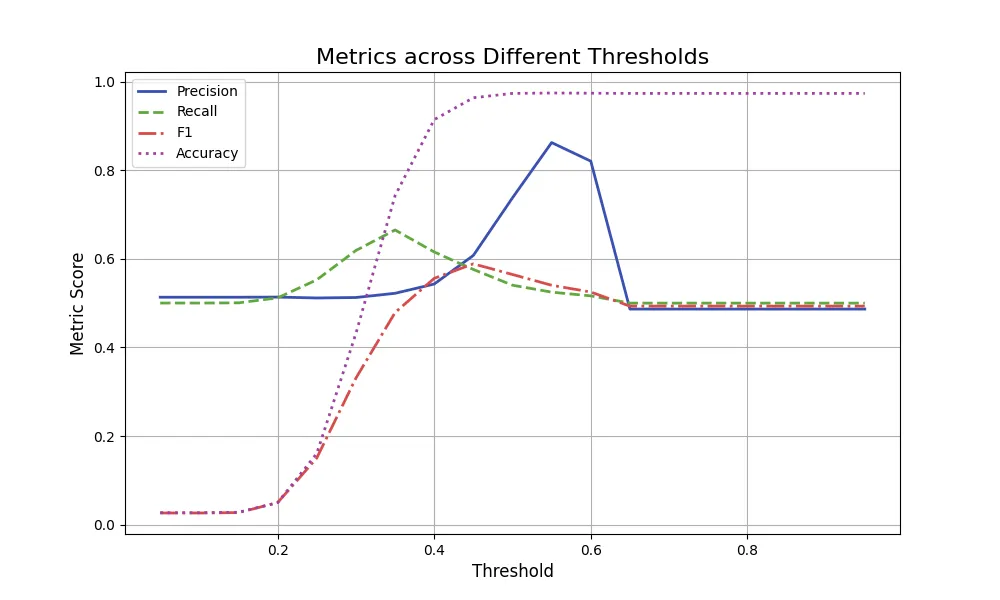

***Supplementary Figure 7:*** *Performance of the sex-specific BirdNET classifier across different confidence thresholds for detecting NF male and female songs from a 20% testing dataset (Validation). Precision, recall, F1-score, and accuracy are shown as a function of the confidence threshold.*

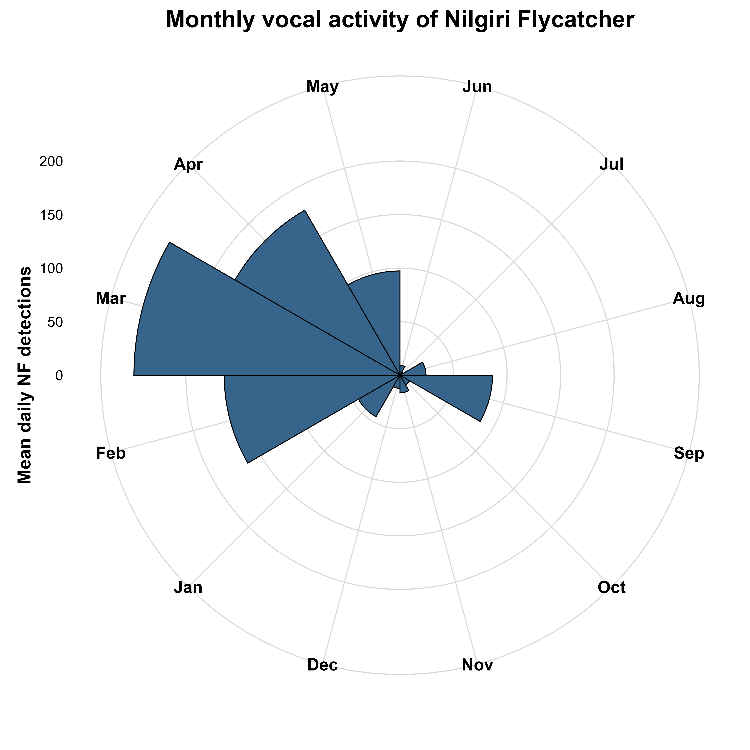

***Supplementary Figure 8:*** *Radial plot showing the monthly variation in the mean daily BirdNET detections of the Nilgiri flycatcher vocal activity from June 2024 to May 2025.*
